# Identifying chemogenetic interactions from CRISPR knockout screens with drugZ

**DOI:** 10.1101/232736

**Authors:** Medina Colic, Gang Wang, Michal Zimmermann, Keith Mascall, Megan McLaughlin, Lori Bertolet, W. Frank Lenoir, Jason Moffat, Stephane Angers, Daniel Durocher, Traver Hart

## Abstract

Chemogenetic profiling enables the identification of gene mutations that enhance or suppress the activity of chemical compounds. This knowledge provides insights into drug mechanism-of-action, genetic vulnerabilities, and resistance mechanisms, all of which may help stratify patient populations and improve drug efficacy. CRISPR-based screening enables sensitive detection of drug-gene interactions directly in human cells, but until recently has largely been used to screen only for resistance mechanisms. We present drugZ, an algorithm for identifying both synergistic and suppressor chemogenetic interactions from CRISPR screens. DrugZ identifies synthetic lethal interactions between PARP inhibitors and both known and novel members of the DNA damage repair pathway. Additionally, drugZ confirms KEAP1 loss as a resistance factor for ERK inhibitors in oncogenic KRAS backgrounds and identifies additional cell-specific mechanisms of drug resistance. The software is available at github.com/hart-lab/drugz.

## Introduction

The ability to systematically interrogate multiple genetic backgrounds with chemical perturbagens is known as chemogenetic profiling. While this approach has many applications in chemical biology, it is particularly relevant to cancer therapy, where clinical compounds or chemical probes are profiled to identify mutations that inform on genetic vulnerabilities, resistance mechanisms, or targets [1]. Systematic surveys of the fitness effects of environmental perturbagens across the yeast deletion collection [2] offered insight into gene function at a large scale, while profiling of drug sensitivity in heterozygous deletion strains identified genetic backgrounds that give rise to increased drug sensitivity [3]. Now, with the advent of CRISPR technology and its adaptation to pooled library screens in mammalian cells, high-resolution chemogenetic screens can be carried out directly in human cells [4–6]. Major advantages to this approach include the ability to probe all human genes, not just orthologs of model organisms; the analysis of how drug-gene interactions vary across different tissue types, genetic backgrounds, and epigenetic states; and the identification of suppressor as well as synergistic interactions, that may preemptively indicate mechanisms of acquired resistance or pre-existing sources of resistant cells in heterogeneous tumor populations.

Design and analysis of CRISPR-mediated chemogenetic interaction screens in human cells can be problematic. Positive selection screens identifying genes conferring resistance to cellular perturbations typically have a high signal-to-noise ratio, as only mutants in resistance genes survive. This approach has been used to identify genes conferring resistance to targeted therapeutics, including BRAF and MEK inhibitors, as well as other drugs [5, 7–14]. Conversely, negative selection CRISPR screens require growing perturbed cells over 10 or more doublings to allow sensitive detection of genes whose knockout leads to moderate fitness defects. Adding a drug interaction necessitates dosing at sub-lethal levels to balance between maintaining cell viability over a long timecourse and inducing drug-gene interactions beyond native drug effects. To our knowledge, a study by Zimmerman *et al.* [15] and Wang *et al.* [16] last year, which each used an early version of the software described here, represents the first such efforts in cancer cells.

Several algorithms currently exist for the analysis of drug-gene interaction experiments [17, 18]. Most rely on adapting methods originally developed for the analysis of RNAseq differential expression data, which is typically characterized by relatively high read counts across genes. High read counts enable the statistically robust detection and ranking of differential expression of genes (in RNA-seq) or abundance of guide RNA (gRNA, in CRISPR screens) using approaches such as the negative binomial P-value model, a trend explored thoroughly in [18]. However, low read counts per gRNA are common in CRISPR data, and are a fundamental feature of genes with fitness defects, leading to a severe loss of sensitivity when applied to CRISPR screens for synthetic chemogenetic interactions.

In this study, we describe drugZ, an algorithm for the analysis of CRISPR-mediated chemogenetic interaction screens. We apply the algorithm to identify genes that drive normal cellular resistance to the PARP inhibitor olaparib in three cell lines. We demonstrate the greatly enhanced sensitivity of drugZ over contemporary algorithms by showing how it identifies more hits with higher enrichment for the expected DNA damage response pathway, and further how it identifies both synergistic and suppressor interactions. We further demonstrate the discovery of both synergistic and suppressor interactions in a single experiment with KRAS-mutant pancreatic cancer cell lines treated with an ERK inhibitor, and with several first-line therapeutic compounds screened in RPE1 hTERT-immortalized epithelial cells. We provide all software and data necessary to replicate the analyses presented here in our repository at github.com/hart-lab/drugz.

## Results and Discussion

We created the drugZ algorithm to fill a need for a method to identify chemogenetic interactions in CRISPR knockout screens. In a pooled library CRISPR screen, the relative starting abundance of each gRNA in the pool is usually sampled immediately after infection and selection. To identify genes whose knockout results in a fitness defect (“essential genes”), the cells are grown for several doublings and the relative abundance of gRNA is again sampled by deep sequencing of a PCR product amplified from genomic DNA template. The relative frequency of each gRNA is compared to starting gRNA abundance and genes whose targeting gRNA show consistent dropout are considered essential genes.

In a chemogenetic interaction screen, the readout is different: the relative abundance of gRNA in a treated population is compared to the relative abundance of an untreated population at a matched timepoint (Figure 1A). In this context, an experimental design with paired samples should be particularly powerful, as it removes a major source of variability across replicates.

**Figure 1.**
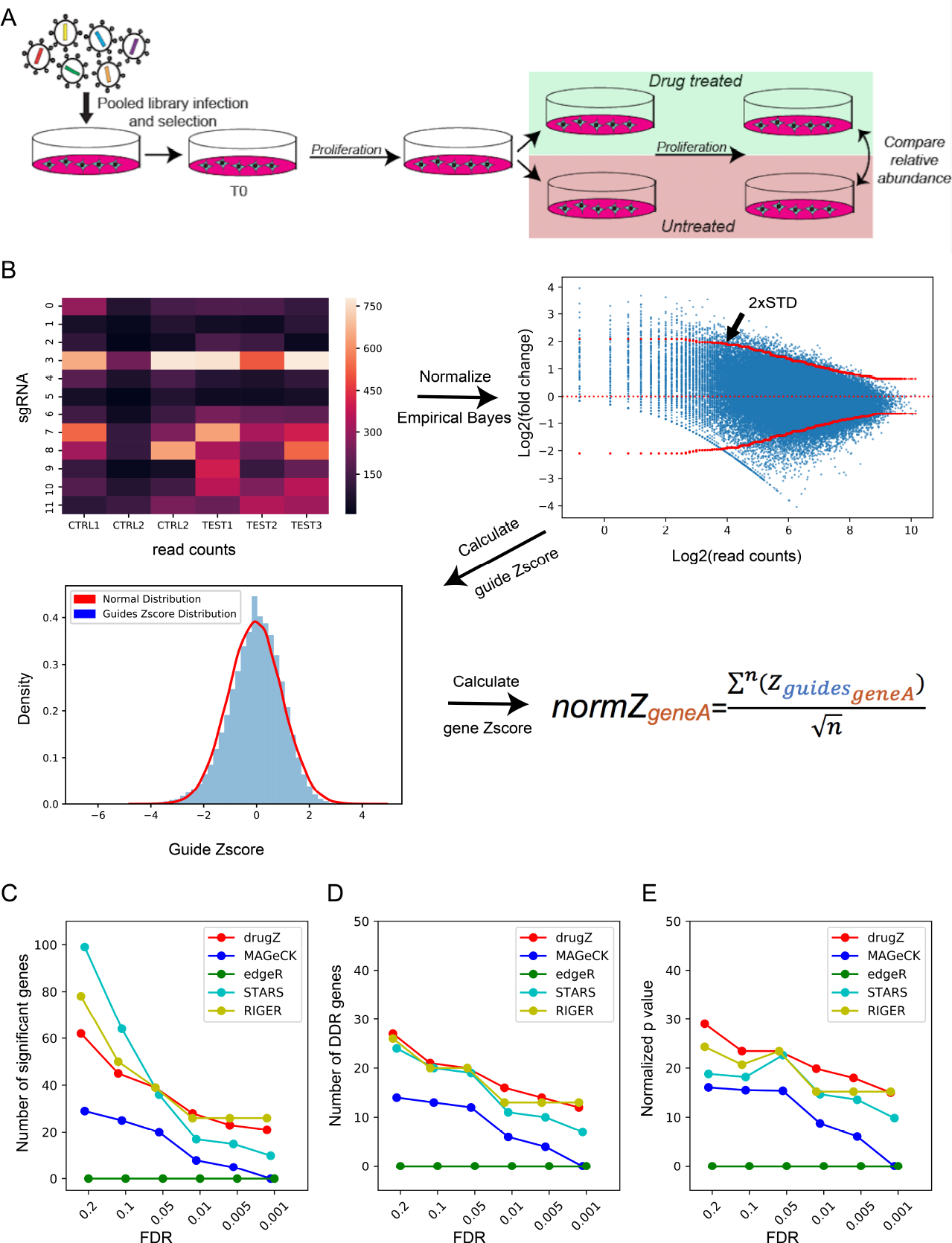
Workflow. **(A)** Experimental design. In a drug-gene interaction screen, cells are transduced with a pooled CRISPR library. Cells are split into drug treated and untreated control samples, grown for several doublings, genomic DNA is collected, and the relative abundance of CRISPR gRNA sequences in the treated and control population is compared. (B) DrugZ processing steps include normalizing read counts, calculating fold change, estimating the standard deviation for each fold change, Z-score transformation, and combing guide scores into a gene score. **(C-E)** Comparing existing methods vs. drugZ for SUM149PT olaparib screen. DrugZ hits show strongest enrichments for DDR genes across a range of FDR thresholds. (C) number of raw hits. (D) number of annotated DNA Damage Response (DDR) genes in hits. (E) -log P-values for DDR gene enrichment by hypergeometric test.

To benchmark the method, we evaluated screens to identify modifiers of the response to the PARP inhibitor olaparib in three cell lines, RPE1-hTERT, HeLa, and SUM149PT [15]. The screens were performed using the TKOv1 library of 90k gRNA targeting 17,000 genes [19]. After infection and selection, each cell line was split into 3 replicates, passaged at least once, and each replicate was further split into control and olaparib-treated populations, providing a paired-sample experimental design (Figure 1A).

The drugZ algorithm calculates a fold change for each gRNA in an experimental condition relative to an untreated control. A Z-score for each fold change is calculated using an empirical Bayes estimate of the standard deviation, by “borrowing” information from gRNA observed at a similar frequency (read count) in the control cells. Guide-level gene scores are combined into a normalized gene-level Z-scores called normZ, from which P-values are estimated from a normal distribution (Figure 1b). We used drugZ to calculate normZ scores, P-values, and false discovery rates in SUM149PT breast cancer cells, which carry BRCA1 and TP53 mutations. We also analyzed the same data with four contemporary methods, STARS [7], MAGeCK [18], edgeR [20], and RIGER [21]. We noted that drugZ produced a moderate number of overall hits, relative to other methods, as FDR thresholds were relaxed (Figure 1c). We evaluated the quality of the hits by measuring their functional coherence. The PARP inhibitor olaparib was developed specifically to exploit the observed synthetic lethal relationship between PARP1 and the BRCA1/BRCA2 genes [22, 23]. Subsequent studies have shown it to be effective against a general deficiency in homologous recombination repair, known as HRD [24]. We therefore calculated the enrichment of each hit set for genes in the DNA damage response (DDR) pathway as annotated in the Reactome database [25] and found that drugZ hits show strong enrichment for DDR genes across a range of FDR thresholds (Figure 1d,e), while the other methods show consistently lower enrichment. We observed similar trends in an olaparib screen in HeLa cells (Supplementary Figure 1A) but less overall effect in RPE1 wildtype epithelial cells (Supplementary Figure 1B). The combination of larger sets of hits and greater enrichment for expected results indicates that drugZ accurately and sensitively identifies chemogenetic interactions.

The drugZ algorithm can also be used to identify suppressor interactions; that is, genes whose perturbation reduces drug efficacy. While *BRCA1* mutation is synthetic lethal with *PARP1*, subsequent mutation of *TP53BP1* is associated with acquired resistance to the PARP inhibitor [26]. Drug-gene interactions resulting in positive Z-scores reflect such suppressor interactions. Indeed, *TP53BP1* is the 8^th^-ranked suppressor interaction in *BRCA1*-deficient SUM149PT cells, with a normZ score of 3.05. Similarly, newly described resistance gene *C20orf196*, now called *SHLD1* [27–30], is the top ranked suppressor.

### Robustness to Parameter Choice and Experimental Design

To evaluate the robustness of the drugZ approach, we conducted sensitivity analysis using data from the SUM149PT olaparib screen. The algorithm relies on two major tunable parameters, window size for empirical Bayes variance estimation and a monotone filter for the variance estimator (to ensure non-decreasing variance as read count decreases). The window size represents the number of neighboring gRNA, ranked by read count, to use to evaluate gRNA fold change variance. To evaluate the effect of varying window size, we ran the drugZ pipeline with window sizes in five increment from 100 to 1,000; neither number of hits, number of DDR-annotated hits, nor enrichment p-value were affected by changing window size (Supplementary Figure 2a). We performed a similar analysis with and without enforcing the monotone filter and discovered marginally improved performance in the SUM149PT olaparib screen without enforcing monotonicity (Supplementary Figure 2b), but no such effect in Hela (T15) olaparib screen (Supplementary Figure 2c). We therefore left the filter in place.

**Figure 2.**
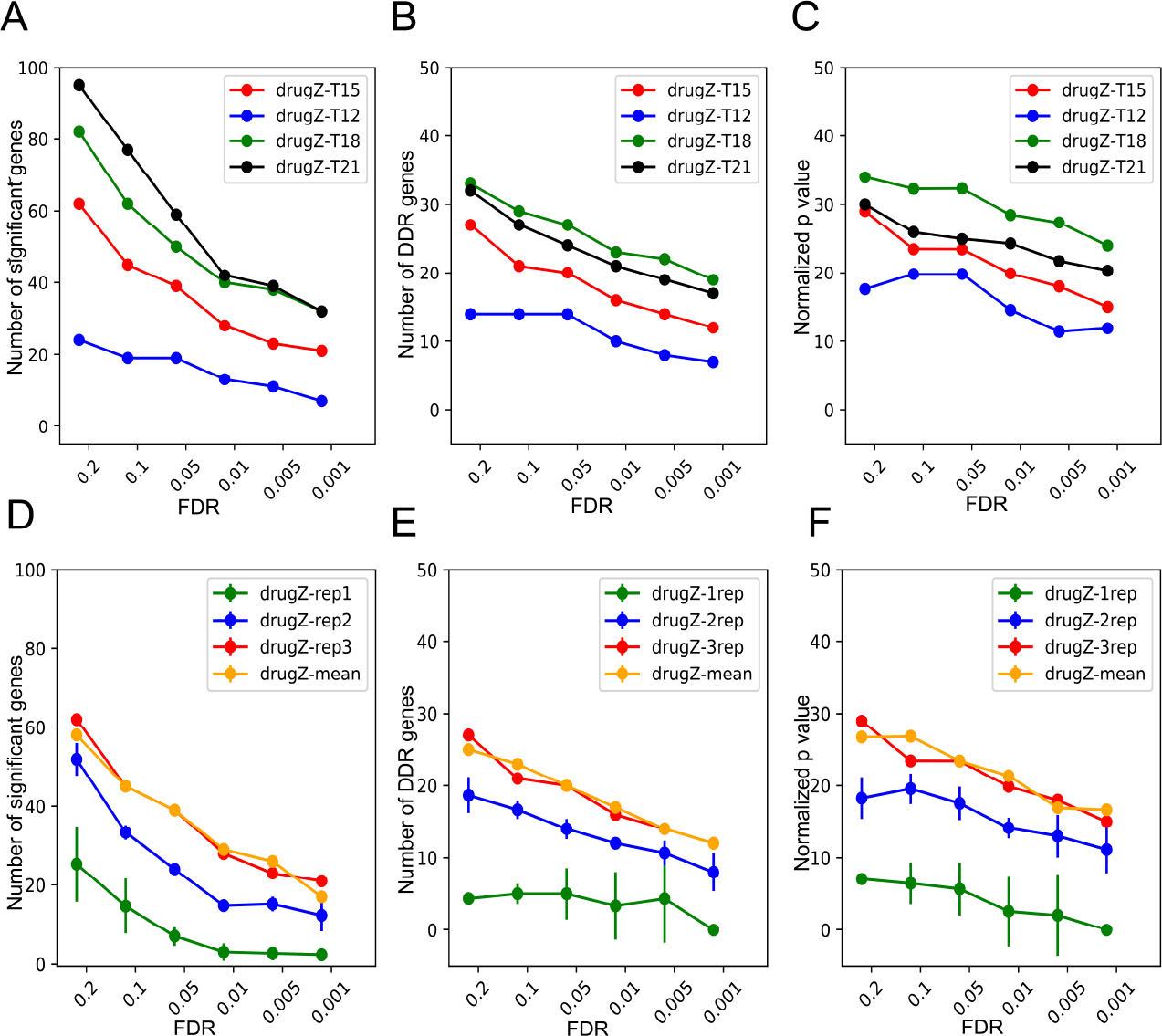
Experimental design effects. **(A-C)** DrugZ performance across different time points for SUM149PT olaparib screen. (A) number of raw hits. (B) number of annotated DNA Damage Response (DDR) genes in hits. (C) -log P-values for DDR gene enrichment. (D-F) DrugZ performance based on varying number of replicates. (D) number of raw hits. (E) number of annotated DNA Damage Response (DDR) genes in hits. (F) -log P-values for DDR gene enrichment. Repl,2,3: all combinations of one, two, or three replicates, +/-s.d. Mean: comparing mean of drug-treated samples to the mean of control samples (unpaired approach).

We also tested the drugZ pipeline against a more statistically thorough, but computationally demanding, approach. After using the same empirical Bayes approach to calculate a Z-score for each guide, we applied Gibbs sampling to estimate the posterior distribution of fold changes for each gene. This method, which we termed drugGS, yielded results that are virtually identical to drugZ (rho=0.99; Supplementary Figure 3B) at ~50x the computational cost (Supplementary Figure 3C). DrugGS is also available on github at https://github.com/hart-lab/druggs.

**Figure 3.**
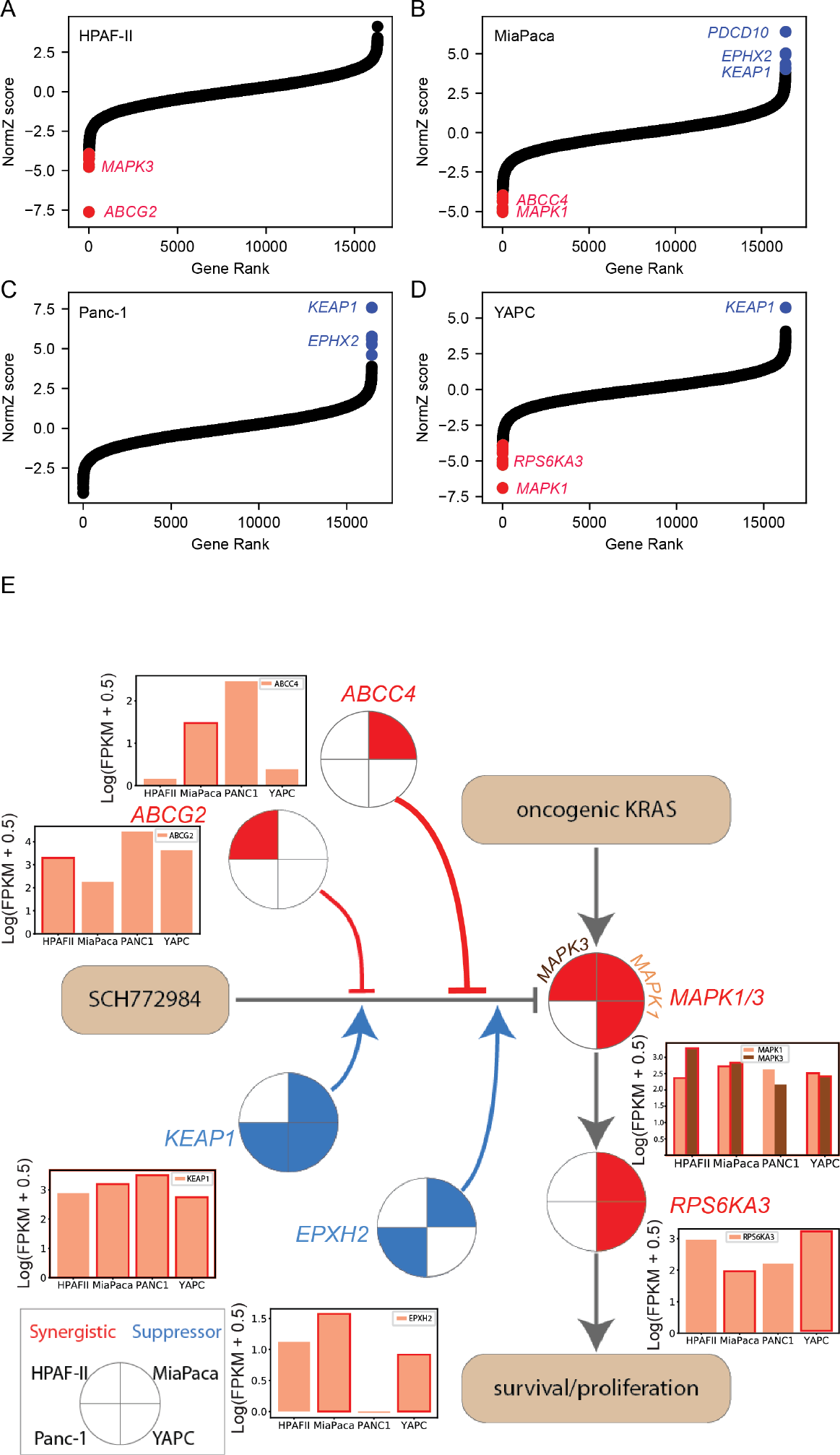
ERK inhibitor screens in pancreatic cancer cell lines. **(A)** drugZ NormZ score is plotted vs. gene rank for SCH772984 screen in HPAF-II pancreatic cancer cells. Red, synergistic (synthetic lethal) interactions at FDR<0.1. (B) MiaPaca cells. Blue, suppressor (resistance) interactions at FDR<0.1. (C) Panc-1 cells. (D) YAPC cells. (E) Network view of ERK inhibitor screens. Red, synthetic lethal interactions. Blue, suppressor interactions. Insets, gene expression of target genes across the four cell lines.

### Experimental design considerations

Highly effective CRISPR knockout screens are done with a variety of experimental designs, with varying numbers of replicates, degree of library coverage, determination of endpoint, and whether intermediate timepoints are included [5–7, 19, 31–37]. The olaparib drug-gene interaction screens described here were performed in triplicate in 15cm plates and passaged every three days, with drug added at day 6 and samples collected for sequencing at each passage starting at day 12. Using the optimized drugZ pipeline, we evaluated each timepoint in the SUM149PT screens. The screen’s ability to resolve specific DNA damage response genes increased steadily from day 12 to day 18 (Figure 2a-c), highlighting the importance of low-dose drug treatment (e.g. LD20). The extended timeframe for the experiment allows greater resolution of negative selection hits as they disappear from the population over several doublings.

Nevertheless, the screens are still quite noisy, necessitating several replicates for accurate assessment of drug-gene interactions. Paired-sample analysis of three replicates in the olaparib screen clearly outperforms one-or two-replicate designs (Figure 2d-f). Surprisingly, however, the paired-sample approach does not appear to offer significant benefits over an unpaired approach: when calculating fold change as the log ratio of the means of three experimental and three control samples, the results are nearly identical to analysis of three paired samples (Figure 2d-f). Indeed, treating samples as paired or unpaired produced highly correlated results (rho>= 0.96) in all three olaparib screens (Supplementary Figure 4a-d).

### A general-use algorithm for drug-gene interactions

To ensure that the drugZ algorithm is not overspecialized for the strong chemogenetic profile of PARP inhibitors, we applied it to a separate set of drug interaction screens in pancreatic cancer cell lines using the ERK1/2 inhibitor SCH772984. Oncogenic mutations in *KRAS* drive constitutive signaling in the MAP kinase pathway and are associated with proliferation and survival signals. Consistent with current models of *RAS* pathway activation, knockout of inhibitor target *MAPK1* and its downstream target *RPS6KA3* have strong synthetic sick/lethal or negative interactions with ERK inhibitor in two of the cell lines, MiaPaca and YAPC (FDR < 0.1; Figure 3a). In the third cell line, HPAF-II, the top synthetic interactors were drug transporter *ABCG2* and *MAPK3*. Activity of this drug resistance gene may account for this cell line’s resistance to ERK inhibition and the lack of other synthetic effectors in this screen. Drug transporter *ABCC4* is synthetic lethal in MiaPaca cells, suggesting multiple context-dependent routes of drug resistance for this molecule. Epoxide hydrolase *EPHX2* and ubiquitin ligase adapter *KEAP1* are the top two suppressors of ERK inhibitor activity in three cell lines, suggesting these genes are required for normal function of the inhibitor (Figure 3b). *KEAP1* loss-of-function was identified as a modulator of MAP kinase pathway inhibitors in a panel of positive selection screens in multiple cell lines[11], but *EPHX2* is a novel candidate resistance gene. Notably, the ERK inhibitor screens yielded a small number of discrete synthetic and suppressor hits, in contrast with the PARP inhibitor screens, which showed broad interaction across the HR pathway, confirming the general applicability of drugZ in detecting drug-gene interactions.

We further tested genetic response profile of hTERT-immortalized RPE1 epithelial cells to two commonly used chemotherapeutic drugs, gemcitabine and vincristine, plus HDAC-inhibitor entinostat currently in clinical trials (Figure 4a). We used our BAGEL pipeline to identify genes whose knockout leads to fitness defect (essential genes; Bayes Factor > 10) or enhanced growth (tumor suppressors, BF < -40) in untreated control cells (Figure 4b). Each drug reveals synthetic lethal interaction with at least one pathway-specific gene. Entinostat, ostensibly an inhibitor of histone deacetylases *HDAC1* and *HDAC3*, is synthetic lethal with *HDAC7* in RPE1 cells. Gemcitabine, a pyrimidine nucleoside analog, is synthetic lethal with deoxythymidylate kinase *DTYMK*. *DTYMK* phosphorylates dTMP to dTDP, a key step in the synthesis-by-salvage pathway of dTTP [38]. Vincristine, a microtubule stabilizer, is synthetic lethal with CLASP1, a nonmotor microtubule-associated protein that promotes kinetochore-microtubule attachment [39]. Vincristine is further synthetic lethal with drug transporter ABCC1 (multidrug resistance protein MRP1), a known marker of vincristine resistance [40, 41].

**Figure 4.**
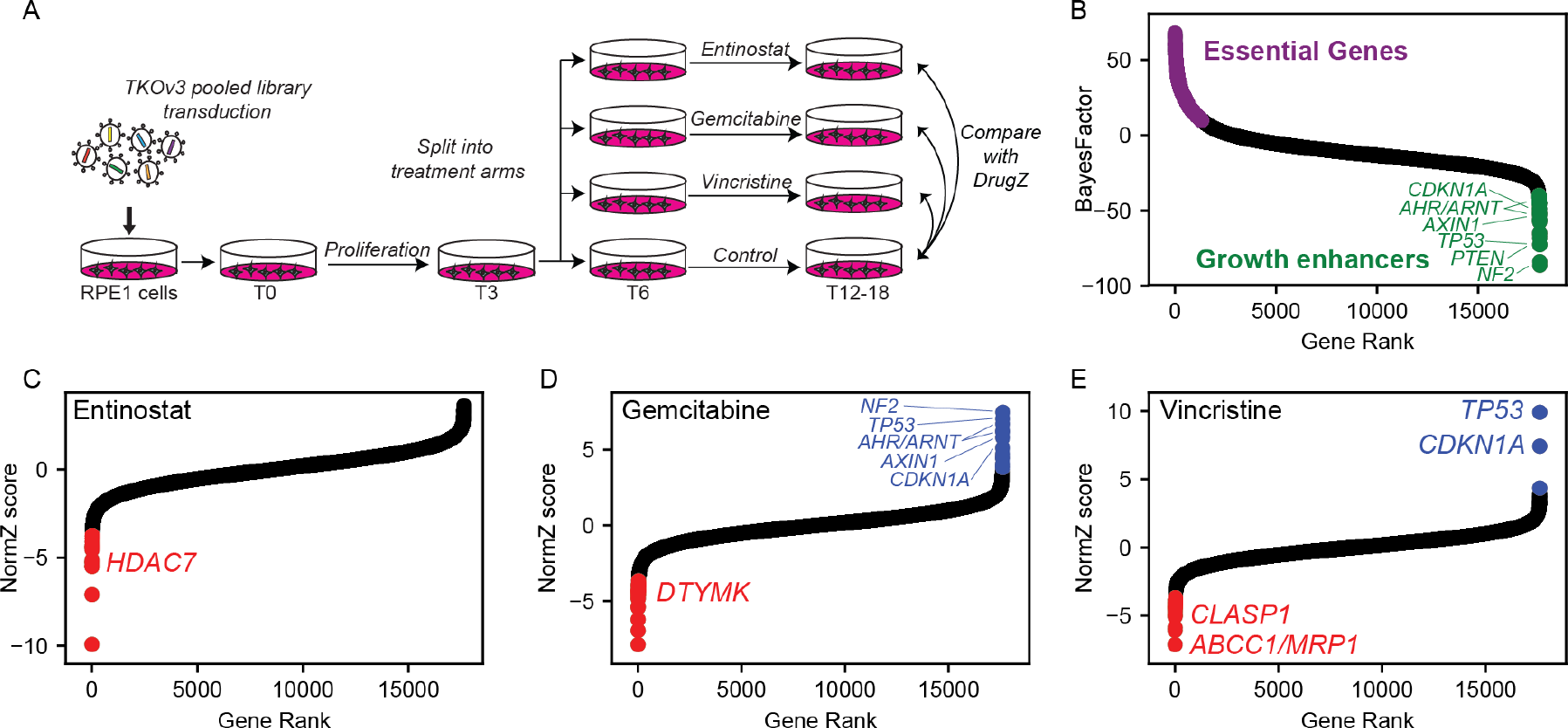
Multiple drug screens in hTERT-RPE1 cells. (A) Experimental design. The TKOv3 lentivirial library was transduced into RPE1 cells, expanded, and split into four treatment arms (in duplicate). (B) Control cells were analyzed with BAGEL to identify essential genes (purple) and putative tumor suppressors (green). (C) NormZ scores for RPE1 entinostat screen; colors as in Figure 3. (D) NormZ scores for RPE1 gemcitabine screen. (E) NormZ scores for RPE1 vincristine screen.

Two of the three drugs appear to show suppressor interactions with known tumor suppressors, including TP53/CDKN1A, NF2, and the aryl hydrocarbon receptor complex AHR/ARNT. This epistatic interaction is probably driven by the drug treatment masking the growth-enhancing effect of knocking out these genes rather than a clinically useful drug-gene interaction. The growth-enhancing effects of knocking out tumor suppressor genes in responsive cell lines is likely to be a systematic source of false positives for suppressor interactions using this approach.

## Conclusions

Identifying the genetic drivers of drug effectiveness and resistance is critical to realize the promise of personalized medicine. Chemogenetic interaction screens in mammalian cells using CRISPR knockout libraries have so far been primarily used in a positive selection format to identify the genes, pathways and mechanisms of acquired resistance to chemotherapeutic drugs. However, negative selection screens to identify the underlying architecture of drug-gene interactions have been difficult to carry out and to analyze in part due to the lack of robust analytical tools.

We describe the drugZ algorithm, which calculates a gene-level Z-score for pooled library CRISPR drug-gene interaction screens. By taking into account the moderate single mutant fitness defects associated with many genes involved in drug-gene interactions, the drugZ algorithm offers significantly improved sensitivity over contemporary analysis platforms. The algorithm was developed to exploit the additional resolving power we expected to gain from a paired-sample experimental design, but surprisingly this has virtually no effect on our results. We demonstrate the validity of our hits by showing the strong enrichment for genes involved in the DNA damage response in a screen for interactions with the PARP inhibitor olaparib and the precise detection of MAPK pathway effectors in an ERK inhibitor screen. We further show that both synergistic and suppressor interactions can be identified in the same screen, as the previously identified PARP resistance gene *TP53BP1* and newly characterized *SHLD1* (formerly *C20orf196*) are top-ranked suppressors of olaparib activity in *BRCA1*-mutant SUM149PT screens. Moreover, both synthetic targets *MAPK1/3* and *RPS6KA3* and suppressor genes *EPHX2* and *KEAP1* are identified in ERK inhibition screens. *KEAP1* deletion or mutation is frequently found in KRAS-driven lung adenocarcinomas and may present an obstacle to ERK inhibitor therapy in these tumors.

Experimental design plays a critical role in the ability to accurately identify drug-gene interactions. Negative selection screens for synthetic lethal interactions require that cells be carried long enough for dropouts – typically growth defects rather than full synthetic lethals – to rise to statistical significance. Our results, concordant with known highly drug-specific differences in effect timing, suggest that each there is value in collecting multiple timepoints to ensure that drug activity and genetic interaction are detectable, and that traditional dose-response curves must be calculated over a timecourse relevant to the screen (e.g. at least two passages or several doublings).

Despite these technical idiosyncrasies, chemogenetic interaction screens extend the utility of CRISPR genome-scale perturbation screens by enabling the systematic surveying of the landscape of drug-gene interactions across cancer-relevant genetic backgrounds. Understanding this variation may lead to more precise therapies for patients as well as the development of synergistic drug combinations for genotype-specific treatments.

## Acknowledgments

MC, GW, WFL, and TH were supported by MD Anderson Cancer Center Support Grant P30 CA016672 (the Bioinformatics Shared Resource) and the Cancer Prevention Research Institute of Texas (CPRIT) grant RR160032, and TH is supported by NIGMS grant R35GM130119. MZ is a Banting postdoctoral fellow. Work in the DD lab was funded through CIHR grant FDN143343, Canadian Cancer Society grant #70389, as well as a Grant-in-Aid from the Krembil Foundation. Work in JM lab is funded through CIHR grants 342551 and 365646. Work in SA lab was funded through a CIHR grant #361837.

## Data and material availability

All software described in this manuscript, as well as all data files used for analysis, are available at the Hart Lab github site (github.com/hart-lab/drugz) as well as the Hart Lab website (hart-lab.org).

## Methods

### DrugZ algorithm

We calculate the log_2_ fold change of each gRNA in the pool by normalizing the total read count of each sample (to n=10 million reads) at the same timepoint and taking the log ratio, for each replicate, of treated to control reads.

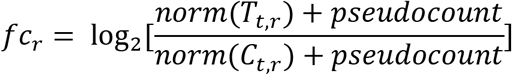

Where:

- fc = fold change
- r = replicate indication
- T = treated sample
- C = control sample
- t = time point
- pseudocount = default value is 5

We estimate the variance of each fold change by calculating the standard deviation of fold changes with similar abundance in the control sample:

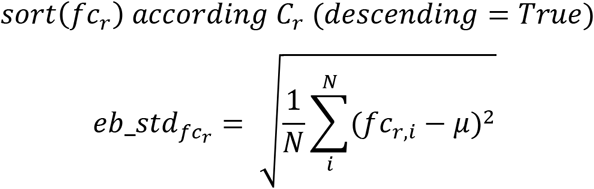

Where:

- *eb_std*_*fc*_*r*__ = estimated variance
- *N* = number of fold changes with similar abundance (default = 1000)
- *i* = guide
- *fc*_*r,i*_ = fold change for each guide in a replicate
- *μ* = 0

and then calculate a Z-score for each fold change using this estimate:

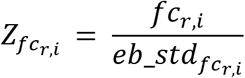

The guide Z score of all gRNA across all replicates is summed to get a gene-level sumZ score, which is then normalized (by dividing by the square root of the number of summed terms) to the final normZ (Figure 1B)

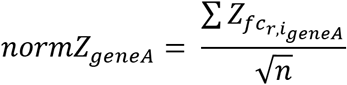

A P-value is calculated from the normZ, and corrected for multiple hypothesis testing using the method of Benjamini and Hochberg [42]. The open-source Python software can be downloaded from github.com/hart-lab/drugz.

### DrugGS algorithm

After Empirical Bayes variance estimation approach is applied on normalized log fold changes to calculate a Z-score for each guide, we applied Gibbs sampling to generate posterior distribution of fold changes for each gene.

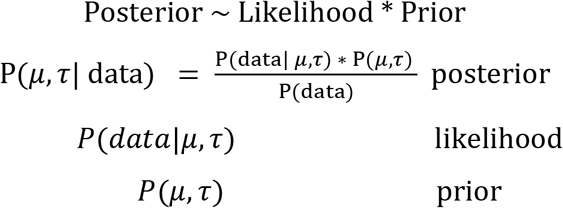

Each gene has a distribution composed of Z-scores for guides targeting that specific gene across replicates. Distribution is characterized as ℕ(μ, τ), where *τ* is 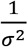.

Both *μ and τ* have hyperparameters (*μ*: *μ*, *σ*^2^, *τ*: *a*, *b*) that we initialize at the very start of sampling.

P(*τ*| data) ~ Γ(a, b) = Gamma prior with a (shape) and b (rate) hyperparameters

P(*μ*|*τ*, data) ~ ℕ(*μ*, *σ*^2^) = Normal prior with *μ* (mean) and *σ*^2^ (variance) hyperparameters

We then update *μ and τ* with respect to their priors in every of 1000 samples that we generate for each gene.

Equations to update *μ*:

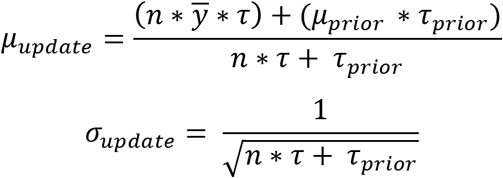

Equations to update *τ*:

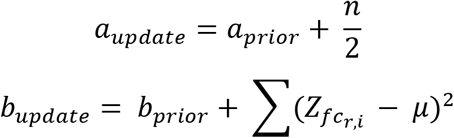

Where:

- n = number of data points (guide Z scores) for each gene
- 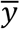 = actual mean of data points

From those 1000 newly sampled *μ and τ*, we then calculate mean and standard deviation. Each gene’s *μ* posterior distribution’s mean is what was converted into Z score and used to compare with the drugZ normZ values.

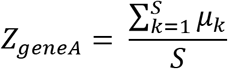

Where:

- S = number of samples (in our case 1000)
- k = sample

### Drug-Gene interaction screens

Olaparib screens were described in [15].

### Cell Culture

hTERT RPE-1 (CRL-4000) and 293T (CRL-3216) cells were purchased from the ATCC and grown in Dulbecco’s High Glucose Modified Eagle Medium (DMEM;HyClone) with 10% fetal bovine serum (FBS), 1 X GlutaMAX (Gibco), 100mM sodium pyruvate (Gibco), 1 X non-essential amino acids (NEAA), 1X penicillin-streptomycin (Pen/Strep), and 5ug ml^−1^ Plasmocure. Incubator conditions were kept at 37°C with 5% CO_2_.

### Lentivirus Production

For production of the TKOV3 lentivirus, 9.0 X 10^6^ 293T cells were transfected with psPAX2 (lentiviral packaging; Addgene #12260), pMD2.G (VSV-G envelope; Addgene #12259), and TKOV3 (Toronto KnockOut CRISPR Library; Addgene #90294) using X-tremeGENE 9 DNA transfection reagent (Sigma-Aldrich) in medium with lowered antibiotic concentration (0.1X Pen/Strep). Medium was replaced with viral harvest medium (DMEM + 1.1% BSA + 1X Pen/Strep) 18 hours post-trasfection. Virus-containing supernatant was collected ~24-48 hours post-transfection, and fresh viral harvest medium was added to transfected plates. Virus-containing supernatant was collected again ~24 later. The virus-containing supernatant was centrifuged to remove cell debris and stored at -80°C.

### CRISPR screening

For transduction of the hTERT RPE-1 cells, the TKOv3 virus was added with 8ug/ml Polybrene. For selection of the transduced cells, puromycin was introduced at a concentration of 20 ug/ml at 24 hours post-infection (the hTERT cassette used to immortalize RPE1 cells contains a puromycin resistance marker, necessitating extreme puromycin concentrations for selection). Puromycin selection continued for 72 hours post-transduction and completed upon the selection against the hTERT RPE-1 parental line as a control. Completion of selection was considered the initial timepoint (T_0_). The TKOv3-transduced cells were split into technical replicates. To ensure proper coverage, 15 x 10^6^ cells across 11 x 15 cm dishes were used for infection with the TKOv3 virus per replicate. The chemotherapeutic drugs Entinostat (2nM), Gemcitabine (2nM), and Vincristine (0.4nM) were added to separate replicates, with one set of replicates receiving no drug treatment. Both drug-treated and untreated replicates were not allowed to reach confluence in the 15cm dishes. Cells were lifted, counted, and re-plated at the coverage stated above, and the excess cell pellets were frozen at -20°C as a timepoint. Once 8 doublings were reached from T_0_, the screens were terminated and pellets frozen at -20°C. Coverage of screens was kept at 200 cells per gRNA.

The QIAamp Blood Maxi Kit (Qiagen) was used to isolate the genomic DNA (gDNA) from the frozen cell pellets. Guide sequences were enriched using PCR with HiFi HotStart ReadyMix (Kapa Biosystems) and primers targeting the guide region in the genomic DNA. A second round of PCR was performed with i5 and i7 primers to give each condition and replicate a unique multiplexing barcode. The final PCR products were purified using the E-Gel System (Invitrogen), normalized, and sequenced on the NextSeq500 system to determine the representation of guides under each treated and non-treated condition.

---

**Supplementary Figure 1.**
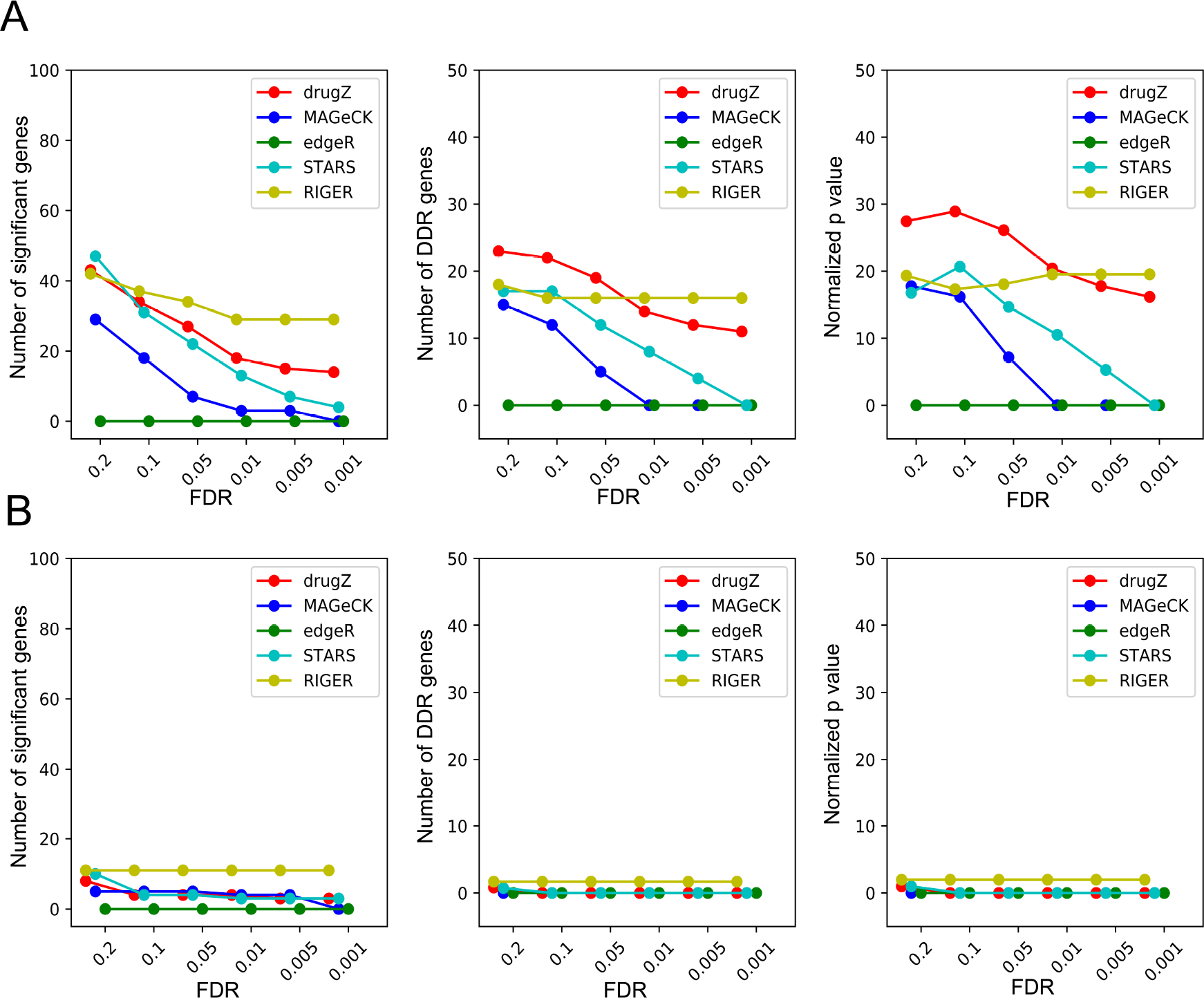
DrugZ vs. other methods for HeLa (A) and RPE1 (B) with olaparib screens. Methods are colored as in Fig.1C. DrugZ hits show strongest enrichment for DDR genes across a range of FDR thresholds in these two screens as well but less overall effect in RPE1 cells. **(A)** Left, number of raw hits. Center, number of annotated DNA Damage Response (DDR) genes in hits. Right, log P-values for DDR gene enrichment. **(B)** All three panels are the same as in (A), for RPE1 screen.

**Supplementary Figure 2.**
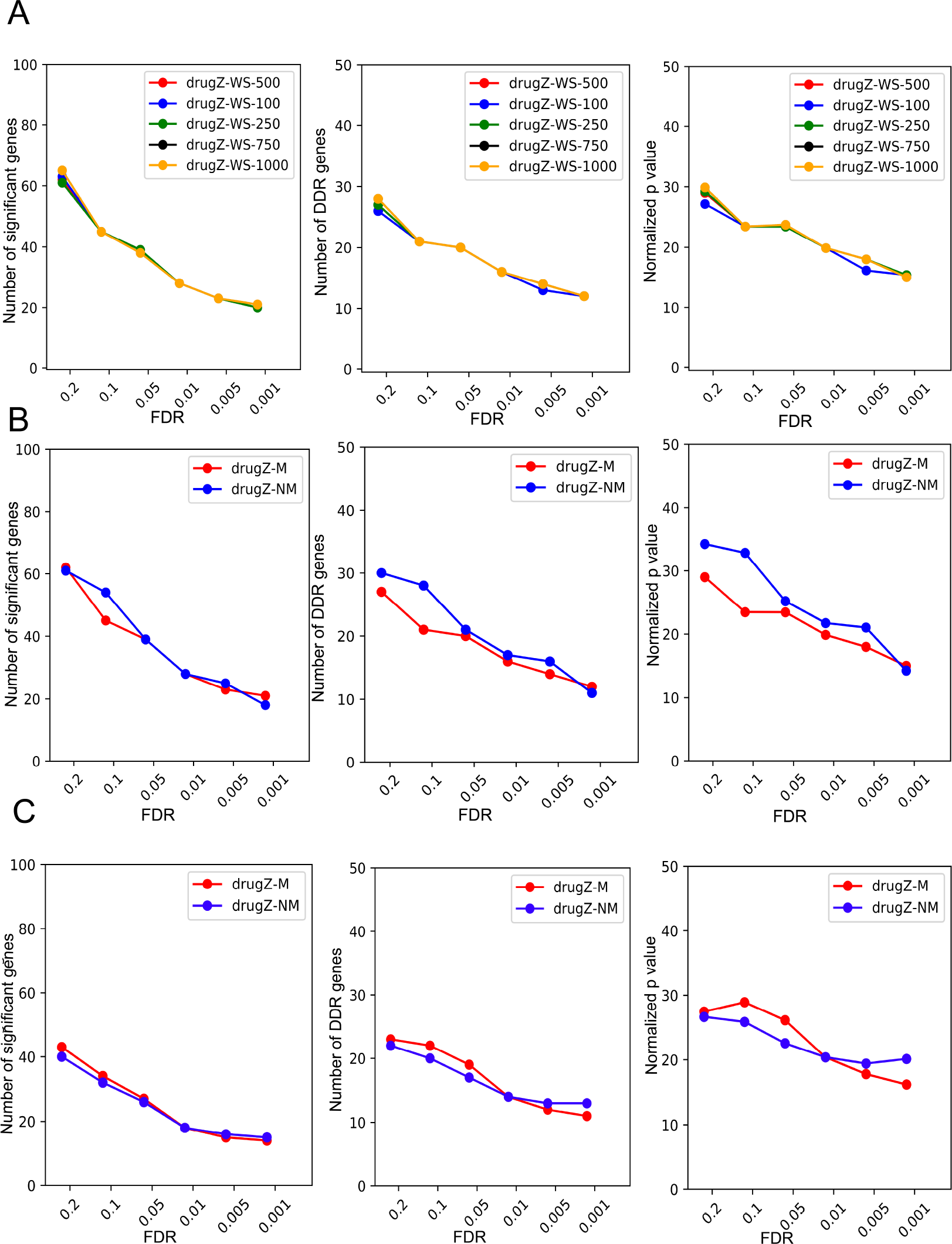
DrugZ tunable parameters. **(A)** DrugZ performance across different window sizes for Empirical Bayes estimation of variance. Left, number of raw hits. Center, number of annotated DNA Damage Response (DDR) genes in hits. Right, log P-values for DDR gene enrichment. **(B)** DrugZ performance with correction that ensures monotonicity in the variance (red) vs. drugZ performance with no correction that ensures monotonicity in the variance (blue) in SUM149PT olaparib screen (panels same as in (A)). **(C)** DrugZ performance with correction that ensures monotonicity in the variance (red) vs. drugZ performance with no correction that ensures monotonicity in the variance (blue) in HeLa olaparib screen (panels same as in (A) and (B)).

**Supplementary Figure 3.**
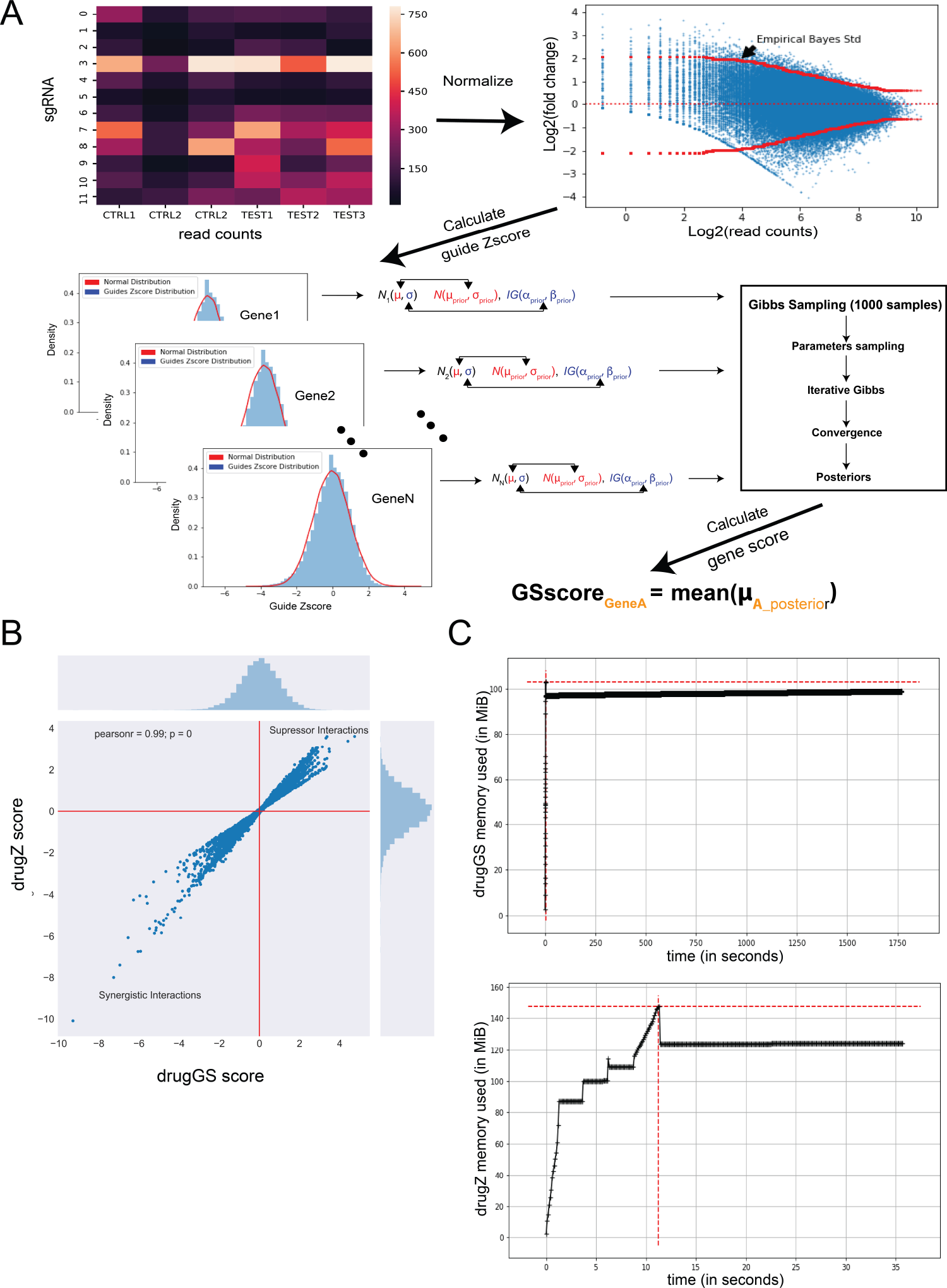
DrugZ vs. DrugGS. **(A)** DrugGS Computational Diagram. DrugGS processing step are same as in the DrugZ until the step where the gene-level scores are generated. After guide level Z-scores are obtained, they are used as a prior distribution for gene-level score in Gibbs sampler. The mean of generated samples of means is considered as new gene score. **(B)** Comparison between drugGS (x-axis) and drugZ (y-axis) gene scores. High correlation between the two (rho = 0.99). **(C)** Comparison between drugGS (top) and drugZ (bottom) time and memory performance. DrugZ drastically outperforms drugGS in terms of time and memory used.

**Supplementary Figure 4.**
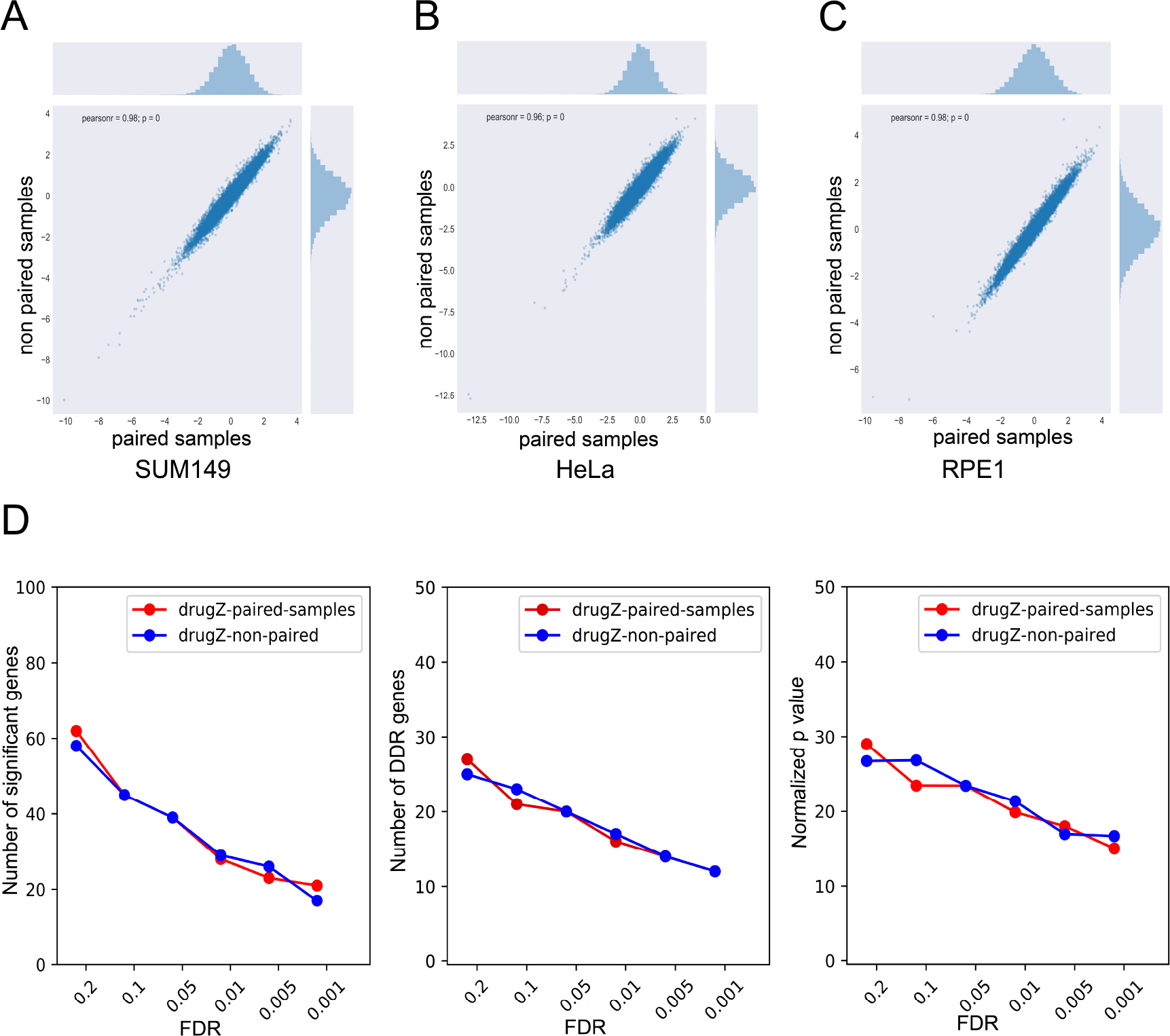
High correlation between paired and non-paired approaches in there olaparib screens. **(A)** Correlation between paired samples (control A – treated A, control B – treated B, etc.) vs. non-paired (mean (control A, B, C) – mean (drug A, B, C.)) for Sum149 olaparib screen (rho = 0.98) **(B)** Same as in (A) for HeLa olaparib screen (rho = 0.96) **(C)** Same as in (A) for RPE1 olaparib screen (rho = 0.98) **(D)** Comparison between paired and nor-paired approaches across number of significant genes, DDR genes and normalized p-values in SUM149PT olaparib screen.

